# An in-silico study to determine whether changes to mitochondria organization through engineered mitochondrial dynamics can enhance bioenergetics in cardiomyocytes

**DOI:** 10.1101/2020.09.21.307306

**Authors:** Adarsh Kumbhari, Shouryadipta Ghosh, Peter S. Kim, Vijay Rajagopal

## Abstract

Mitochondria are the powerhouse of the cell and owing to their unique energetic demands, heart muscles contain a high density of mitochondria. In conditions such as heart failure and diabetes-induced heart disease, changes in the organization of cardiac mitochondria are common. While recent studies have also shown that cardiac mitochondria split and fuse throughout the cell, a mechanistic understanding of how mitochondrial dynamics may affect energy output is lacking. Using a mathematical model that has been fitted to experimental data, we test if briefly altering fission or fusion rates improves ATP production and supply in cardiomyocytes. Unexpectedly, we found that cardiac bioenergetics, e.g., the ADP/ATP ratio, were robust to changes in fusion and fission rates and consequently mitochondria organization. Our study highlights complex nonlinear feedback loops that are at play in the cross-talk between mitochondrial dynamics and bioenergetics. The study motivate further in-silico and experimental investigations to determine the mechanistic basis for new therapies that target mitochondrial dynamics.

## INTRODUCTION

Mitochondria meet cellular energy demands by converting nutrients into chemical energy in the form of adenosine triphosphate (ATP). In cardiomyocytes, mitochondria have a particularly challenging job, as the heart must pump up to six liters of blood per minute, continuously from birth (Cattermole et al., 2017). Far from being static, mitochondria are continually splitting and fusing in response to energy demands and stressors (Youle and van der Bliek, 2012). While there is an emerging body of experimental work on fission and fusion, there are no quantitative frameworks that investigate if changes to mitochondria arrangement through fission and fusion, can help with bioenergetics.

Cardiac mitochondria are organized into networks of clusters that are interspersed between contractile protein bundles (Ghosh et al., 2018; Glancy et al., 2017). Furthermore, recent studies have shown that cardiac mitochondria dynamically change their organization (Ong et al., 2015; Glancy et al., 2017) and that it is disrupted in diseased cells (Cao and Zheng, 2019; Galloway and Yoon, 2015). For example, Chen et al. (2012) observed that deficiencies in mitochondrial fusion proteins Mfn1 and Mfn2 cause mild cardiomyopathy. Additionally, defects in the mitochondrial fusion protein OPA1 may cause left ventricular hypertrophy (Piquereau et al., 2012), leading to an increased risk of arrhythmias and heart failure (Frey et al., 2004; Frey and Olson, 2003). Studies by Jarosz et al. (2017) and others have also shown altered organization of mitochondria in diabetes-induced heart disease.

A key idea is that mitochondrial fission and fusion ensure mitochondrial quality control. For example, mitochondrial fusion mediates internal machinery sharing, such as mitochondrial respiration and the equilibration of mitochondrial membrane potential (Chen et al., 2005; Eisner et al., 2014; Eisner et al., 2018; Twig et al., 2008). Thus, by diluting mitochondrial damage, fusion facilitates quality control. Drp1 governs mitochondrial fission (Eisner et al., 2018; Ong et al., 2015), which facilitates quality control by promoting the fragmentation of highly damaged mitochondria and limits the propagation of mitochondrial dysfunction (Eisner et al., 2018; Glancy et al., 2017; Ong et al., 2015). Fission decreases during periods of bioenergetic stress (Gomes et al., 2011; Liesa and Shirihai, 2013), but may increase locally to minimize the spread of mitochondrial dysfunction.

Additionally, mitochondria may increase their mass via biogenesis in response to nutrient deprivation or biophysical stress (Jager et al., 2007; Mihaylova and Shaw, 2011; Scarpulla, 2011). Mitochondrial biogenesis is mediated by the protein PGC-1α and facilitates quality control by promoting mitochondrial homeostasis (Boland et al., 2013; Dalmasso et al., 2017). Despite tantalizing evidence that mitochondrial dynamics could be a possible therapeutic target to improve mitochondrial function and hence energy production (Ong et al., 2015; Ong et al., 2010), the precise mechanistic and causal relationship between mitochondrial dynamics and bioenergetics is still to be explored and would help identify effective drug targets that are based on the underlying mechanism.

We have recently shown that changes to mitochondria morphology and their spatial distribution can influence the distribution of energy metabolites and consequently contractile force under hypoxic, and high-workload conditions (Jarosz et al., 2017; Ghosh et al., 2018). These findings were based on a biophysics-based computational model of mitochondrial function and spatially detailed geometric models of cardiomyocyte architecture derived from electron microscopy images. The works of Eisner et al. (2017) and Glancy et al. (2017) suggest that mitochondrial fusion and fission rates increase at high workloads. These experimental and computational studies suggest that dynamic alterations to mitochondria density and organization, via altered fusion and fission rates, might help meet high energy demands at high workloads.

In this study, we integrated the current understanding, that is outlined above, of the role of fusion/fission dynamics, biogenesis, and mitochondria organization on cardiac bioenergetics into a semi-quantitative mechanistic modelling framework. We have considered three plausible mechanisms by which mitochondrial dynamics and biogenesis can regulate mitochondrial bioenergetics: (i) increased mitochondrial connectivity can enhance the OXPHOS capacity of individual mitochondria; (ii) increased mitochondrial volume can increase the total OXPHOS capacity of the cell; and (iii) mitochondrial network reorganization can favorably alter the diffusion distances between mitochondria and myofibrils for a rapid and steady supply of ATP. We used this model to test whether increasing or decreasing fusion or fission rates would affect bioenergetics. Specifically, we sought to: (i) investigate whether mitochondria network morphological changes stemming from alterations in network connectivity affect bioenergetics in physiological and pathological conditions; (ii) investigate how altered bioenergetics could affect fusion/fission dynamics; and (iii) determine key parameters that need to be measured to robustly validate or negate our model predictions and thus formalize a mechanistic model of the link between mitochondrial dynamics and bioenergetics. As a simplifying assumption, we limit our study to investigate cross-talk on an acute scale of two minutes, which suffices to observe dynamic changes in mitochondria network morphology experimentally (Glancy et al., 2017).

Our computational model is a hybrid agent-based- and partial differential equation model. The agent-based model (ABM) simulates changes in mitochondrial connectivity and mitochondrial mass such as fission, fusion, and biogenesis, while the partial differential equation (PDE) system models various bioenergetic interactions such as oxidative-phosphorylation (OXPHOS), ATP hydrolysis, and the breakdown of reactive oxygen species. In particular, we assumed that mitochondrial connectivity, via fission and fusion, directly feeds forward into OXPHOS and electron transport chain (ETC) kinetics, which then feeds back into fission and fusion dynamics. We then calibrated the model against existing experimental data on cardiac mitochondrial bioenergetics and dynamics in the literature.

Remarkably, our simulations show that bioenergetics are robust to varied fission and fusion rates in the short term under physiological conditions. However, fusion and fission may enhance bioenergetics when mitochondrial function is compromised. Since these findings largely depend on ETC enzyme kinetic rates, they highlight the need for experimental measurements of how ETC enzyme kinetics change during mitochondrial fission and fusion. Moreover, we predict that high workloads may increase mitochondrial volume fractions, which may enhance energetics to meet these high workload demands. Indeed, this study reveals a challenging inverse problem, if the ADP/ATP ratio is robust to changes in fission and fusion, when and under what circumstances do fission and fusion impact bioenergetics?

## RESULTS

### A hybrid agent-based model of mitochondrial dynamics and continuum reaction-diffusion model of bioenergetics

Details of the mathematical model equations that were coupled to create our computational model of mitochondrial dynamics and bioenergetics are provided in the Methods section. Figure 1A illustrates the initial geometry used by the model and is inspired by longitudinal views of cardiac cell architecture under the electron microscope. Figure 1B outlines the basic factors that change during interactions. In brief, mitochondrial dynamics and biogenesis are modulated by energetic stress – a catch-all term that encompasses the ratio of ADP-to-ATP, mitochondrial connectivity, and mitochondrial damage. Consequently, changes in energetic stress can alter the mitochondrial network architecture which further leads to alterations in the mitochondrial ATP synthesis rate and the resulting ADP-to-ATP ratio across the cell. The mitochondrial ADP-to-ATP ratio then governs the energetic stress, establishing a feedback loop between mitochondrial dynamics and bioenergetics. Further details are provided in the Methods section, specifically in the subsection titled “Agent based model”.

**Figure 1.**
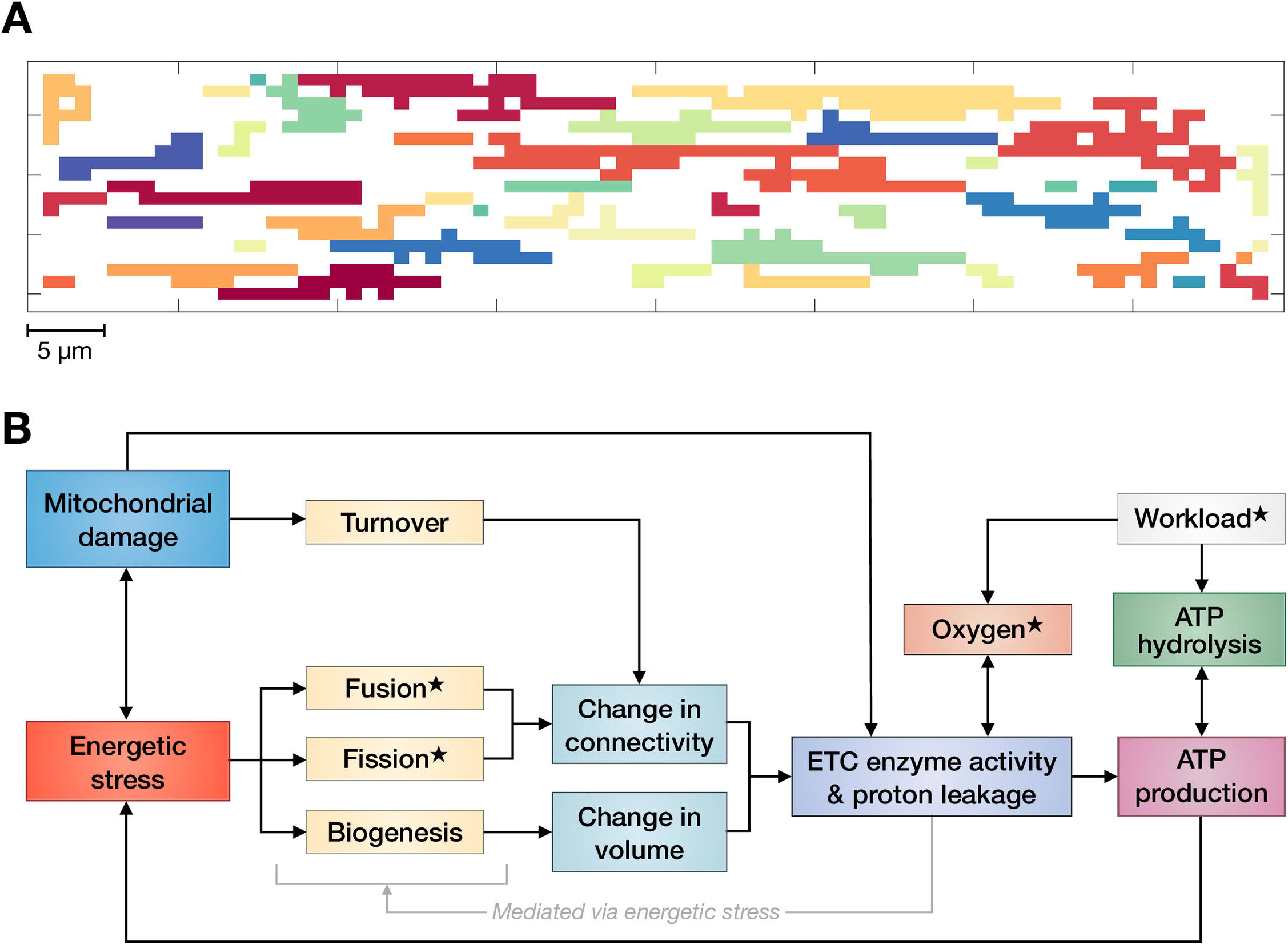
Initial model geometry. (A) Initial conditions used for simulations. Colors denote locally unique mitochondrial clusters. (B) Block diagram depicting the model set up. Starred boxes indicate variables that we vary in this study.

Figure 2 demonstrates that the model captures the key bioenergetic and mitochondrial dynamics features found in current experimental data in the literature. In particular, the model reproduces intracellular ADP (see Figure 2A) and O_2_ distributions (see Figure 2B) similar to those reported by Vendelin et al. (2000) and Takahashi et al. (1998). To simulate the different workloads depicted in Figure 2A, we individually vary *X*_ATPase_, a parameter that quantifies ATP consumption (see Methods, specifically the subsection “ATP consumption”, for specific details). To emulate the experimental set up of Takahashi et al. (1998) in Figure 2B, we change the boundary value of O_2_ from 47.25 μM to 21 μM, which results in a parabolic O_2_ distribution *qualitatively* similar to that reported by Takahashi et al. (1998).

**Figure 2.**
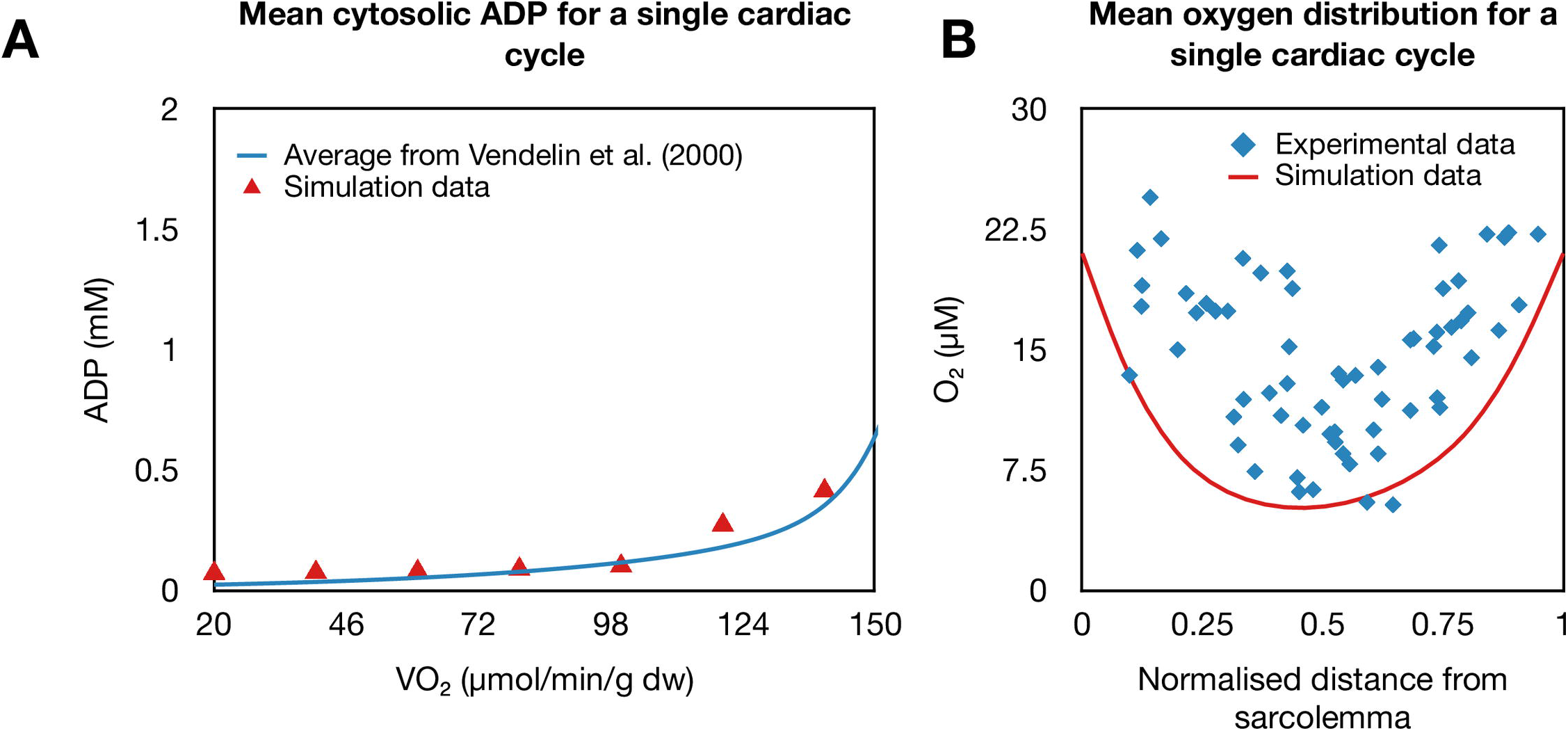
The model reproduces dynamics from the experimental literature. (A) Comparison between our model predictions for spatially averaged ADP vs VO_2_ against data from Vendelin et al. (2000). (B) Comparison between model predictions for radial oxygen profiles against data from Takahashi et al. (1998).

### The model predicts that rate changes in fission and fusion rates do not impact ADP/ATP ratios in healthy cardiomyocytes

To determine the impact of fission and fusion rates on the average ADP/ATP ratio in healthy cells, we simultaneously vary our characteristic fission and fusion rates, λ_split_ and λ_fuse_, over a range of −80% to 200% while holding all other parameters constant at their base values. These characteristic fission and fusion rates are linked to the probability of a fission or fusion event occurring via the following formulas:

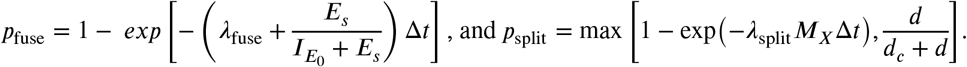

In these formulas, Δ*t* is the size of each ABM time step, *E*_*s*_ describes energetic stress (defined in Equation 44), *M*_*X*_ denotes the mitochondrial mass of a given mitochondrial matrix, and *d* describes mitochondrial damage. Further details are given in the Methods section, specifically the subsection “Agent based model”. Given that *large* changes to either fission or fusion are likely to be highly deleterious (Ong et al., 2015), we assume our range of −80% to 200% is physiologically reasonable. For each fusion or fission rate, we consider the average ADP/ATP ratio from 5 model runs for a simulated duration of 2 minutes. We find that at basal levels, the average ADP/ATP ratio is 9.17×10^−3^ (see Movie S1 for a visualization of the ADP/ATP dynamics predicted by the model). Moreover, we find that varying our fission and fusion rates result in minimal deviations from our basal ADP/ATP ratio despite inducing changes in mitochondria network morphology, specifically, the median mitochondrial cluster size (see Figure 3). This suggests that ADP/ATP ratios in healthy cells are robust to variations in fission and fusion over short timeframes. The same pattern of robustness is also observed in the average PCr/ATP ratio, another bioenergetic parameter that is used to assess cardiac performance.

**Figure 3.**
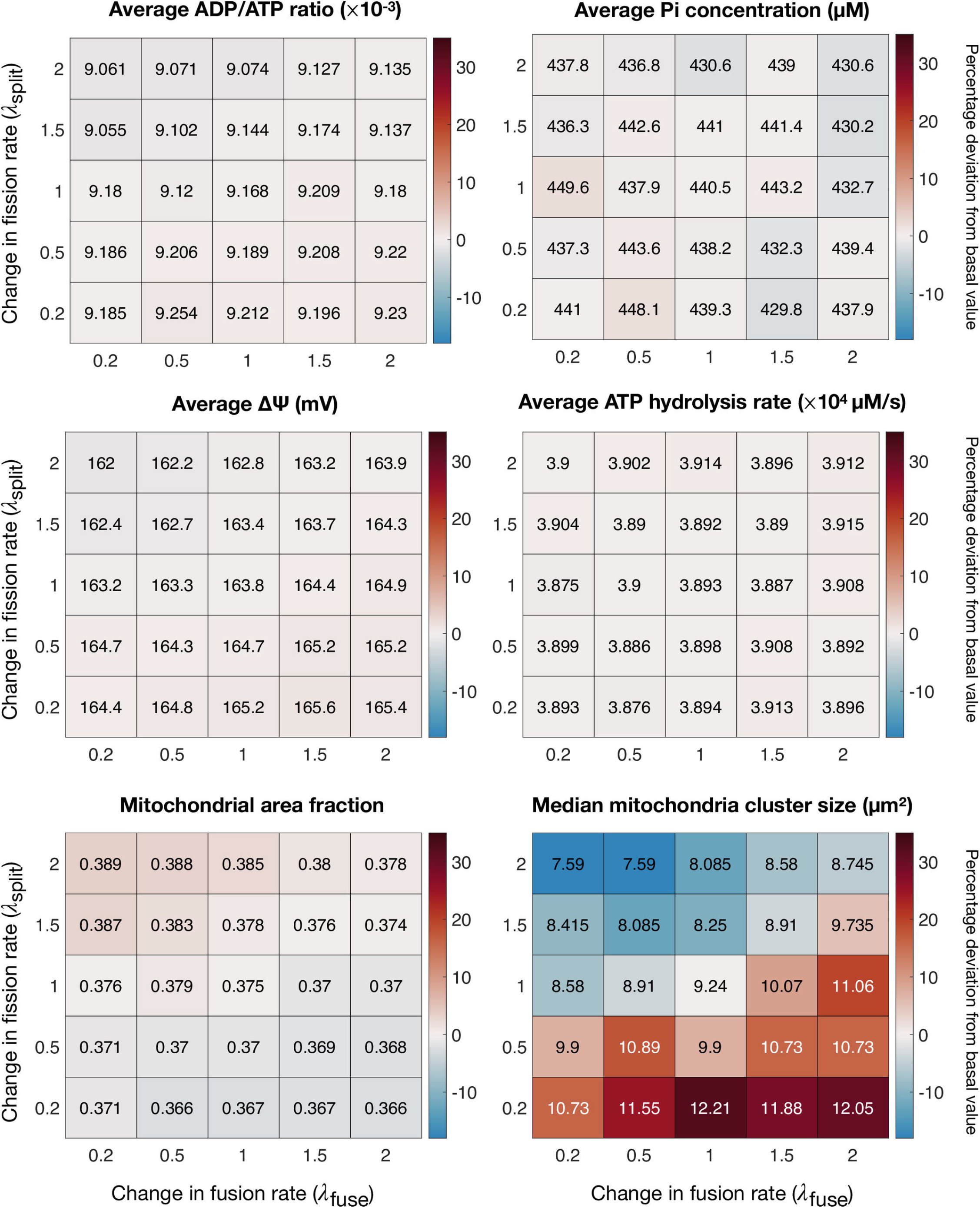
Bioenergetics are robust to rate changes. The average ADP/ATP ratio; average concentration of inorganic phosphate; average membrane potential; average ATP hydrolysis rate; average mitochondrial area fraction; and the median mitochondrial cluster size for varied characteristic fission and fusion rates. Colors denote the percentage deviation from the basal value (no changes to fission and fusion).

To identify the mechanisms that help in maintaining the robustness of ADP/ATP ratios, we analyzed the average mitochondrial membrane potential, the average concentration of inorganic phosphate, and the average ATP hydrolysis rate. We found that the net myofibrillar ATP hydrolysis rate, 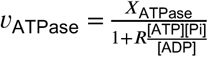 (see Methods, specifically the subsection “ATP consumption”, for more details), which is equivalent to mitochondrial ATP synthesis rate in a steady state, does not vary substantially despite the variation in median mitochondrial cluster size. Similarly, the average membrane potential was also robust to changes in λ_split_ and λ_fuse_.

This bioenergetic stability can be attributed to several mechanisms. In our model, energetic stress modulates the frequency of fission and fusion events. These events alter mitochondria network morphology, which affects OXPHOS activity. As a result, the ADP/ATP ratio changes and with it energetic stress. This change is then integrated into our fission and fusion rates, thus establishing a feedback loop. These feedback controls result in a stable state of mitochondrial dynamics, whereby bioenergetic parameters do not vary substantially, despite changes to the characteristic fission and fusion rates (see Figure 3; also see Movie S1).

In addition to these control mechanisms, robustness is additionally maintained via intracellular shuttling, specifically, adenylate kinase shuttling (Dzeja and Terzic, 2009) and creatine kinase phosphate shuttling (Bessman and Geiger, 1981; Meyer et al., 1984). These shuttles impart an additional layer of robustness to ADP/ATP ratios and ATP hydrolysis rates for varied characteristic fission and fusion rates.

### Model predicts high workloads increase dynamism while hypoxia causes mitochondrial clustering

#### High workloads

To determine how mitochondrial dynamics are acutely affected by an increased workload, we track the number of fission and fusion events that occur for VO_2_ values ranging from 80 μmol min^-1^ g dw^-1^ to 140 μmol min^-1^ g dw^-1^. These particular values are motivated by calculations by Vendelin et al. (2000), who estimate the largest physiological VO_2_ in adult rat hearts to be 160 μmol min^-1^ g dw^-1^. As such, VO_2_ values ranging from 80 μmol min^-1^ g dw^-1^ (50% of the largest physiological value) to 140 μmol min^-1^ g dw^-1^ (87.5% of the largest physiological value), describe high workload conditions. We found that the number of fusion events increased with workload (see Figure 4A) for the entire range of VO_2_. This is a consequence of the gradual rise in ADP/ATP ratio (see Figure 2A) which contributes to an elevation of energetic stress, which in turn increases the rates of fusion and biogenesis. However, higher energetic stress also leads to an increased likelihood of mitochondrial damage (see Equation 48), which would result in a slight increase in fission (see Equation 47) and membrane depolarization (see Figure 4B; see also Equation 31). The net effects of higher fission and fusion rates are larger median sizes of mitochondrial clusters (see Figure 4B) which is consistent with experimental studies (Picard et al., 2013; Yoo et al., 2019). It is important to note that these studies track changes on a scale of hours. By contrast, our simulations track changes on a scale of minutes. As such, increases in the median cluster size may not represent true biogenesis (which occurs on a scale of ∼ 23 minutes at basal conditions, Dalmasso et al. (2017)), but rather aggregation as a result of increases in mitochondrial outer membrane connectivity which can take place within a shorter time frame (scale of seconds) (Glancy et al., 2017). Nevertheless, these findings highlight how, by modulating the frequency of mitochondrial dynamics, mitochondria effectively perform network maintenance ensuring consistent energetics even in high workload conditions.

**Figure 4.**
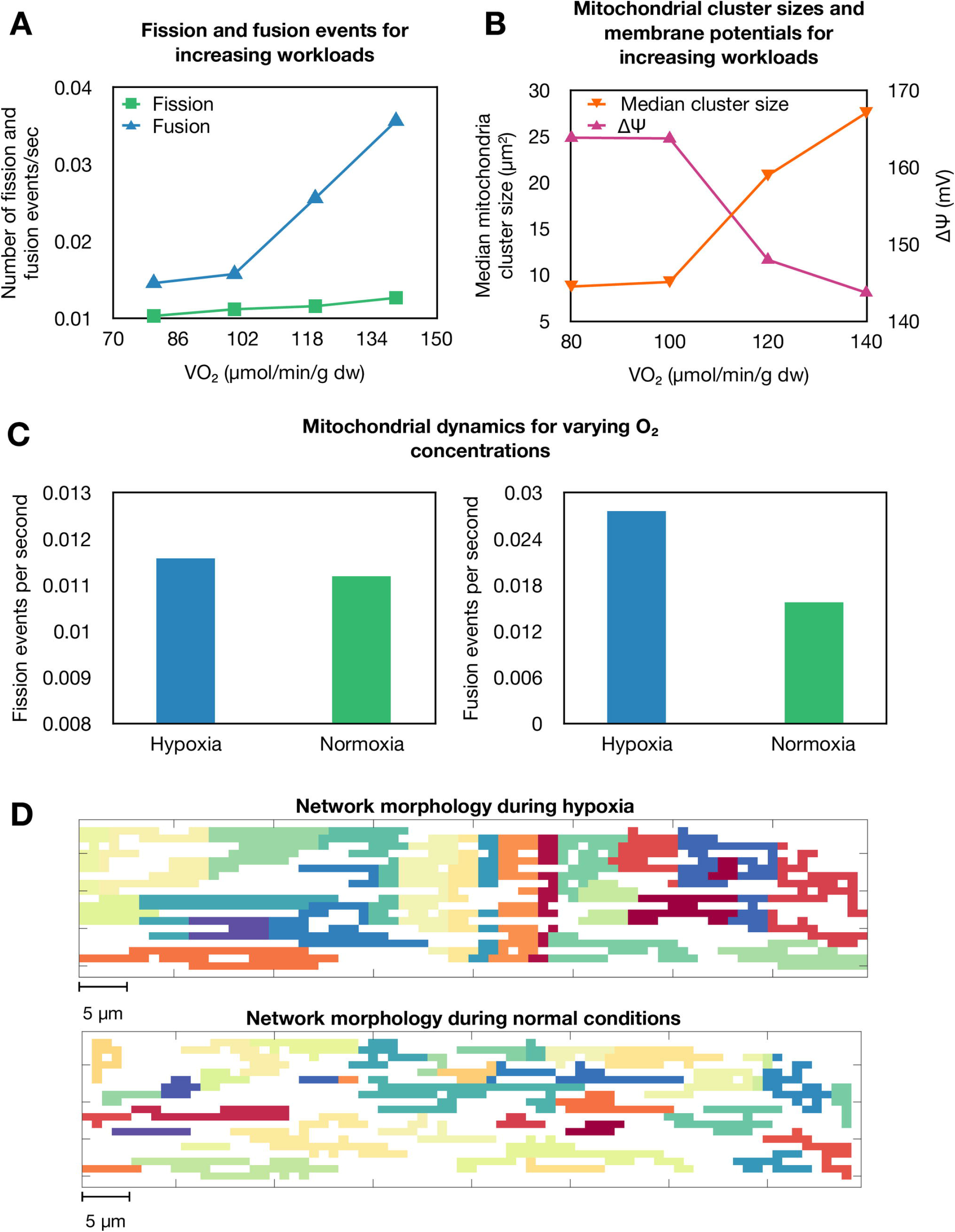
Mitochondrial dynamics under high workloads and hypoxia. (A) Mitochondrial dynamics for various workloads. (B) Median mitochondrial cluster size and mean mitochondrial membrane potential for various workloads. (C) Mitochondrial dynamics in hypoxic conditions. Rates represent averages across the cell for the duration of the simulation. (D) Visualization of the mitochondrial network in hypoxic conditions.

#### Hypoxia

To determine how short-term mitochondrial dynamics differs under hypoxic conditions, we simulate hypoxia and track network fragmentation. Hypoxia is simulated by imposing a constant concentration of 5 μM on the boundary. We found that under hypoxic conditions, mitochondrial membrane potentials were rapidly depolarized (visualized in Movie S2), resulting in an increase in the ADP/ATP ratio (also visualized in Movie S2). Under our modelling assumptions, this increases energetic stress (see Equation 44). Consequently, under hypoxic conditions, mitochondrial networks in our model rapidly increased fusion (see Figure 4C; see also Movie S2). Hypoxic conditions result in certain mitochondrial subnetworks becoming damaged, resulting in an average increase in fission (see Figure 4C) over time. Once segregated, healthy or only mildly damaged mitochondria can fuse to form a robust subnetwork resulting in large mitochondrial clusters (see Figure 4D). This suggests that by segregating damaged subnetworks and fusing together, mitochondria can reduce the spread of dysfunction, thereby allowing the cell to become more robust to hypoxia. The concept of mitochondria acutely segregating damaged subnetworks to improve performance has been also observed in the literature (Glancy et al., 2017).

### Bioenergetics are only mildly robust to altered fission and fusion rates in disease states

Finally, we sought to determine if a disease state, such as diabetes, results in enhanced sensitivity to changes in fission and fusion rates. To answer this, we simulated mitochondrial dysfunction observed in diabetic cardiomyopathy. More specifically, we leveraged a study by Ghosh (2019), in which Beard’s biophysical model of OXPHOS (Beard, 2005) is fit to type I diabetic rat heart data from Pham et al. (2014). That is, to simulate a type I diabetic cell, we decreased the rate of Complex I and Complex V activity by factors of 0.288 and 2.72×10^−4^ respectively and increased the rate of proton leakage by a factor of 1.75. We then simultaneously varied our characteristic fission and fusion rates, λ_split_ and λ_fuse_, over a range of −80% to 200% while simulating a high-intensity workload of VO_2_ = 100 μmol min^-1^ g dw^-1^.

We found that modifications to the rates of fission and fusion still did not markedly improve bioenergetics as mediated by mitochondria network morphology (see Figure 5), despite an increase in the median cluster size. Our simulations suggest that increases in fission, which decrease the median cluster size, are compensated for by an increase in the mitochondrial area fraction. The converse, however, does not appear to be true, i.e., increases in fusion activity do not decrease the mitochondrial area fraction. Importantly, these two compensatory changes in network morphology may help regulate bioenergetics in damaged situations: increased area fractions may enhance bioenergetics by increasing the availability of ATP in the myofibrils (Ghosh, 2019); while increased cluster sizes cause increases in connectivity and thus enhance OXPHOS (as defined in Equation 30, see subsection “ATP production via OXPHOS” within the Methods section for more details). As a result of these feedback mechanisms, the cell maintains an average membrane potential that is robust to changes in fission and fusion. Notably, simulating diabetes does result in the average concentration of Pi being more sensitive (relative to our simulations at basal conditions) to changes in fission and fusion. This is a consequence of Pi regulating mitochondrial metabolism to a greater degree than the ADP/ATP ratio. However, given that type I diabetes is a chronic condition, we accept that on a longer timescale, promoting elongation via fusion – which in our computational model lowers energetic stress – may protect the cell from further damage, either as a result of impaired OXPHOS function or due to external stressors.

**Figure 5.**
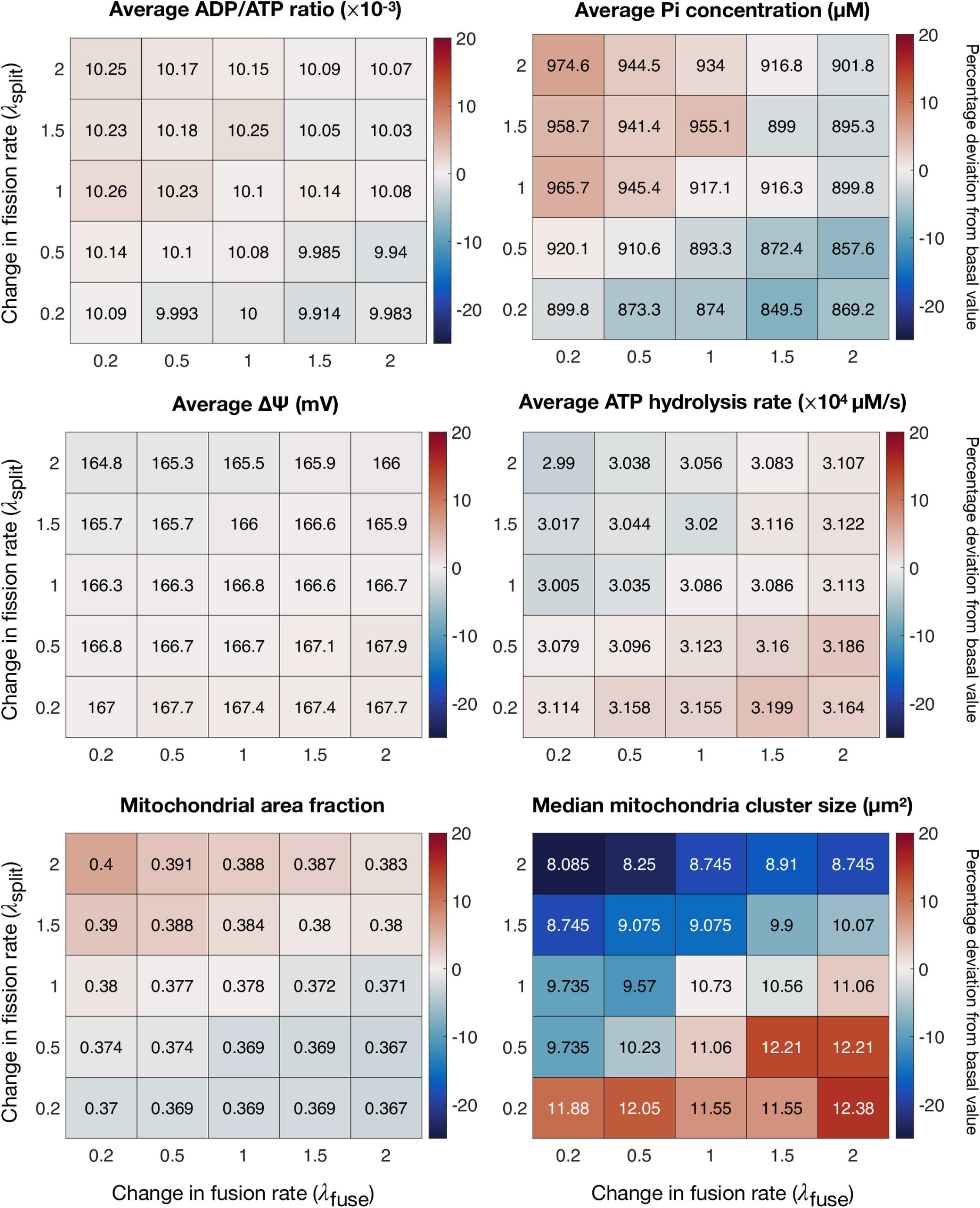
Bioenergetics are mildly robust to rate changes when simulating mitochondrial dysfunction as observed in diabetic cardiomyopathy. The average ADP/ATP ratio; average concentration of inorganic phosphate; average membrane potential; average ATP hydrolysis rate; average mitochondrial area fraction; and the median mitochondrial cluster size for varied characteristic fission and fusion rates. Colors denote the percentage deviation from the basal value (no changes to fission and fusion).

## DISCUSSION

In this study, we develop a semi-mechanistic model to quantitatively explore the range of fission and fusion behaviors that may help with ATP distribution. Our modelling reveals that varied fusion and fusion rates do not result in substantial changes to ADP/ATP ratios in cardiomyocytes in the short term. Furthermore, our modelling shows that changes in connectivity alone do not have an immediate impact on bioenergetics as has been suggested in the literature (Hoitzing et al., 2015).

### Our model highlights the robustness of bioenergetics to changes in mitochondria OXPHOS and fusion/fission properties

Scholars debate the link between ATP synthesis and mitochondrial dynamics. For example, Cipolat et al. (2006); Frezza et al. (2006); Olichon et al. (2003); and Gilkerson et al. (2003), propose that mitochondrial networks can increase ATP production because of changes in membrane shape. Parra et al. (2011) propose that by more uniformly distributing mitochondrial membrane potentials, increased connectivity may improve bioenergetics. In direct contrast, Hoitzing et al. (2015) suggests that mitochondrial dynamics may have no function in relation to increased ATP production. Our model does not involve parameters for individual mitochondrial morphology but identifies two levels of metabolic robustness to changes in fission and fusion rates: mitochondria network morphology and bioenergetics. At the network morphology level, changes in connectivity affect bioenergetics, which controls energetic stress. These changes in stress then modulate the rates of fission and fusion, which establishes a feedback loop, resulting in a stable state of mitochondrial dynamics. At the bioenergetic level, intracellular shuttles such as adenylate kinase shuttling (Dzeja and Terzic, 2009) and creatine phosphate shuttling (Bessman and Geiger, 1981; Meyer et al., 1984) mediate energetic buffering. These feedback mechanisms result in ADP/ATP and PCr/ATP ratios robust to rate changes. Arguably, changing our model of energetic stress to depend on more dynamic factors may reduce this robustness. Future work will address this by using network motifs (Milo et al., 2002; Li et al., 2014), to identify factors that when incorporated into our stress calculation, would increase the sensitivity of ADP/ATP to varied fusion and fusion rates.

### Our model is not a complete representation of the cross-talk between energetics and mitochondrial dynamics

Our model does not account for the pleiotropic effects of fission and fusion on cellular architecture, nor does it account for possible changes in signaling pathways, which may serve as an additional energetic buffer or perhaps even a compensatory role in bioenergetics. For example, in the context of acute ischemic reperfusion injury, Hall et al. (2016) note that inhibiting fusion proteins disrupts the tethering between mitochondria and the sarcoplasmic reticulum, but paradoxically has a cardioprotective effect. Investigating how mitochondrial dynamics reshape cellular architecture is a key area that we will explore in future work. The model representation of mitochondrial networks in two-dimensions and the reduced order representation of individual mitochondrion morphology also remove the possibility to interrogate the role of mitochondrion size and shape on bioenergetics. Nevertheless, the model provides insights on the role that mitochondria fusion/fission dynamics may confer based on current understanding of the relationship between energetics and mitochondrial dynamics.

It is possible that modulation of fusion and fission may indirectly or directly affect mitochondrial expression of respiratory complexes, which could then affect ADP/ATP ratios more drastically. For example, in our model, we assumed that enzyme activity responds linearly to changes in connectivity. However, the transportation rate of metabolites from the IMM to the IMS via ANT saturates out for large ATP concentrations (Beard, 2005), which when coupled with PCr shuttling, maintains a stable ADP/ATP ratio (Bessman and Geiger, 1981; Meyer et al., 1984). Indeed, while dramatic (10^2^ to 10^4^ fold) decreases in enzyme activity, comparable to chronic disease conditions (Wu et al., 2007), do increase the average ADP/ATP ratio across the cell in our simulations, they do not substantially decrease the average mitochondrial membrane potential (see Table 5). Thus, implementing a larger change in enzyme activity due to mitochondrial connectivity is unlikely to *qualitatively* change our findings in the present model.

**Table 1.** List of variables used in PDE model

| Variable | Description |
| --- | --- |
| $t$ | Time (seconds) |
| ATP | ATP concentration ($\mu\text{M}$) |
| MgATP | Mg bound ATP concentration ($\mu\text{M}$) |
| ADP | ADP concentration ($\mu\text{M}$) |
| MgADP | Mg bound ADP concentration ($\mu\text{M}$) |
| AMP | AMP concentration ($\mu\text{M}$) |
| PCr | Phosphocreatine concentration ($\mu\text{M}$) |
| Cr | Creatine concentration ($\mu\text{M}$) |
| Pi | Inorganic phosphate concentration ($\mu\text{M}$) |
| O<sub>2</sub> | Oxygen concentration ($\mu\text{M}$) |
| H<sup>+</sup> | H<sup>+</sup> concentration (also expressed in pH) |
| K<sup>+</sup> | Potassium ion concentration ($\mu\text{M}$) |
| Mg<sup>2+</sup> | Free magnesium ion concentration ($\mu\text{M}$) |
| NADH | NADH concentration ($\mu\text{M}$) |
| NAD | NAD concentration ($\mu\text{M}$) |
| Q | Ubiquinone concentration ($\mu\text{M}$) |
| QH<sub>2</sub> | Ubiquinol concentration ($\mu\text{M}$) |
| Cred | Cytochrome C (reduced) concentration ($\mu\text{M}$) |
| Cox | Cytochrome C (oxidized) concentration ($\mu\text{M}$) |
| $\Delta\Psi$ | Mitochondrial membrane potential (mV) |

**Table 2.** Details of mitochondrial reactions

| Symbol for flux | Description | Source |
| --- | --- | --- |
| $v_{\text{ATPase}}$ | ATP consumption in myofibrils | Wu et al. (2008) |
| $v_{\text{CK}}$ | Creatine kinase reaction | Vendelin et al. (2000) |
| $v_{\text{AK}}$ | Adenylate kinase reaction | Vendelin et al. (2000) |
| $v_{\text{DH}}$ | Dehydrogenase flux representing the TCA cycle and other NADH-producing reactions | Beard (2005) |
| $v_{\text{C1}}, v_{\text{C2}}, v_{\text{C3}}, \text{ and } v_{\text{C5}}$ | Flux through complex I, complex III, complex IV, and complex V ($F_0F_1$ - ATP synthase) | Beard (2005) |
| $v_{\text{leak}}$ | Flux of proton leak across the inner membrane | Beard (2005) |
| $v_{\text{ANT}}$ | Rate of exchange of metabolites through adenine nucleotide translocases (ANT) | Beard (2005) |
| $v_{\text{PiH}}$ | Flux through Phosphate Hydrogen co-transporter | Beard (2005) |
| $v_{\text{KH}}$ | Flux through $K^+ / H^+$ antiporter | Beard (2005) |
| $v_{\text{mtCK}}$ | Flux of mitochondrial creatine kinase reaction | Ghosh (2019) |
| $v_{\text{MiAK}}$ | Flux of mitochondrial adenylate kinase reaction | Vendelin et al. (2000) |

**Table 3.**
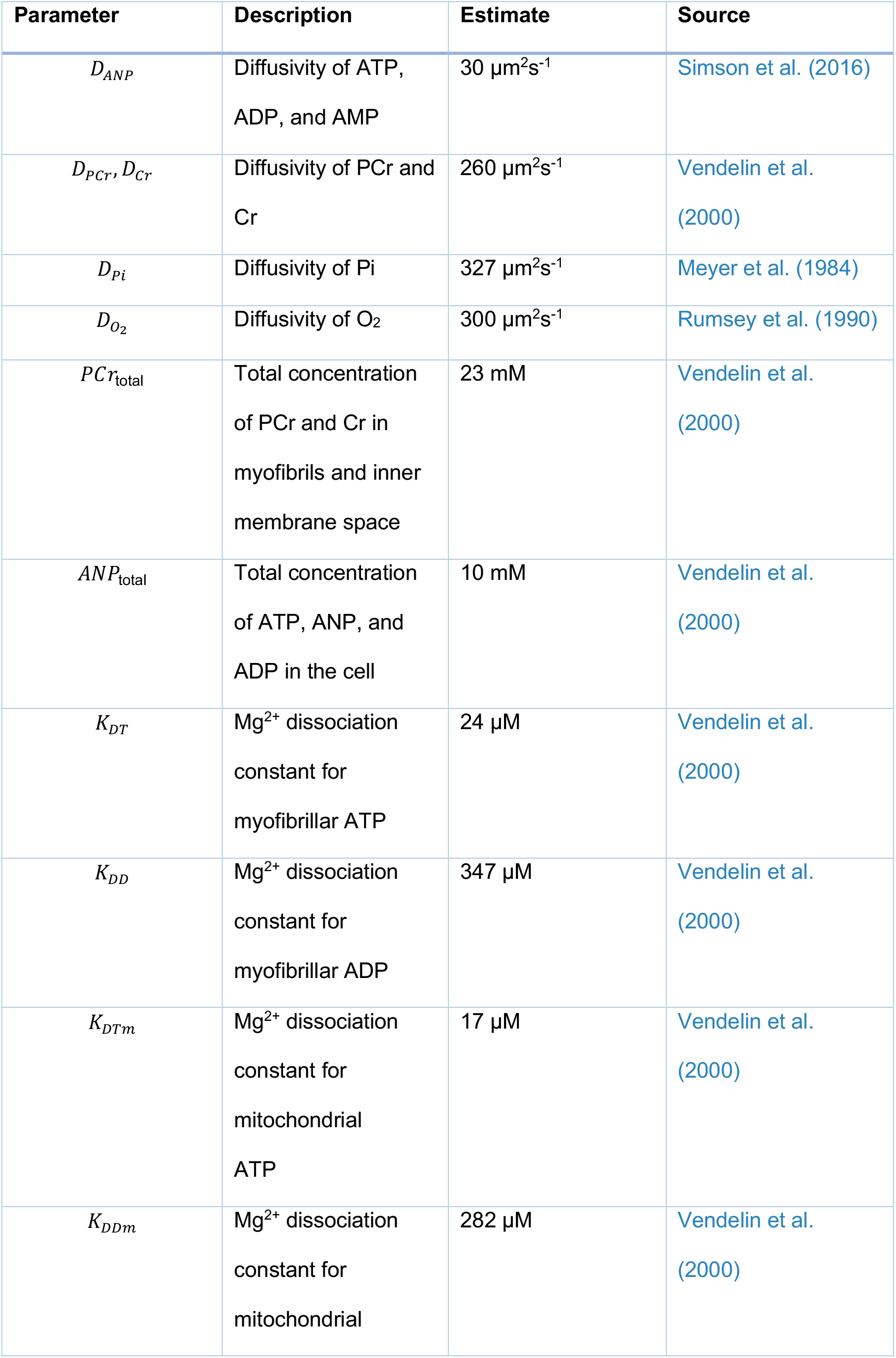
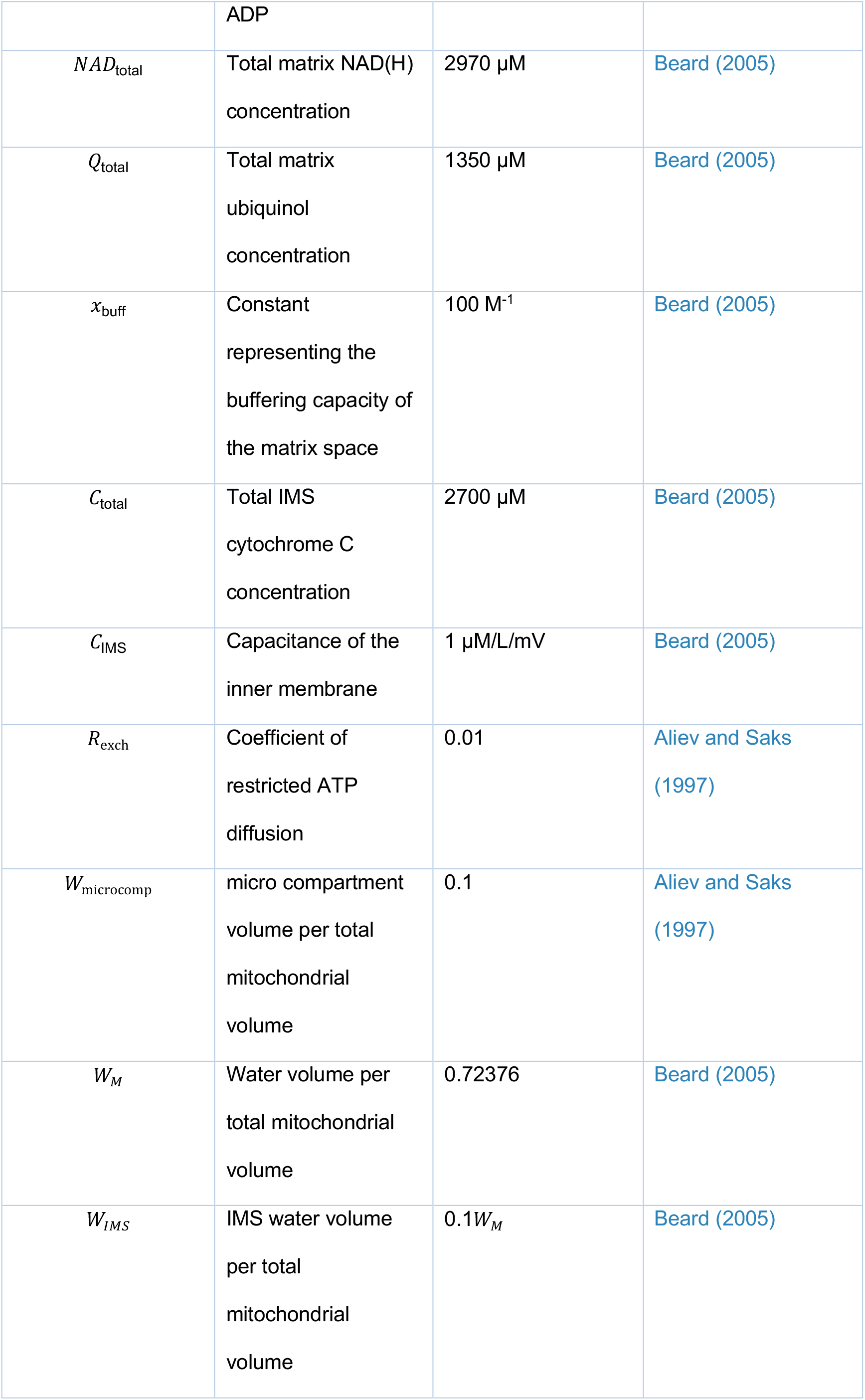
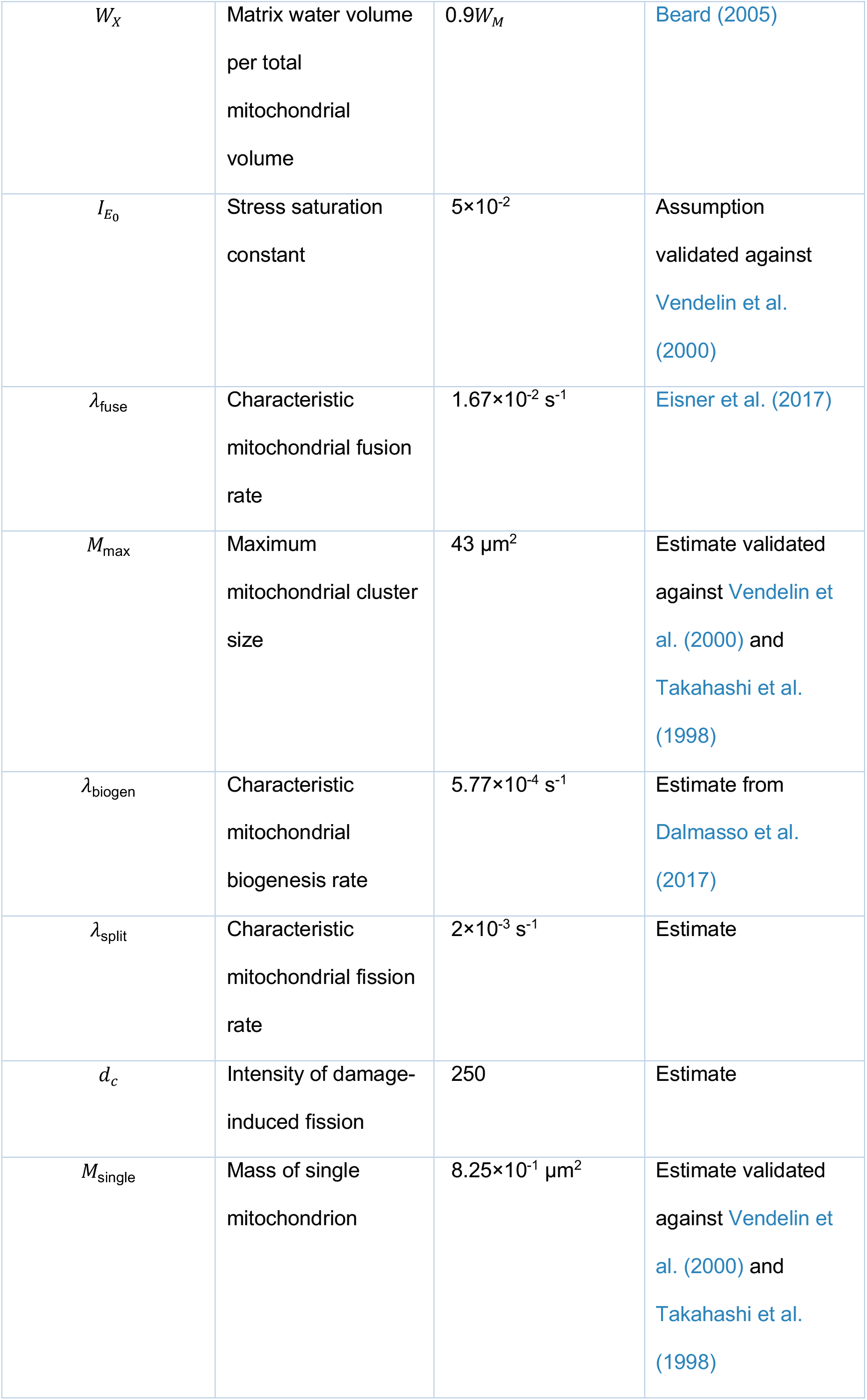
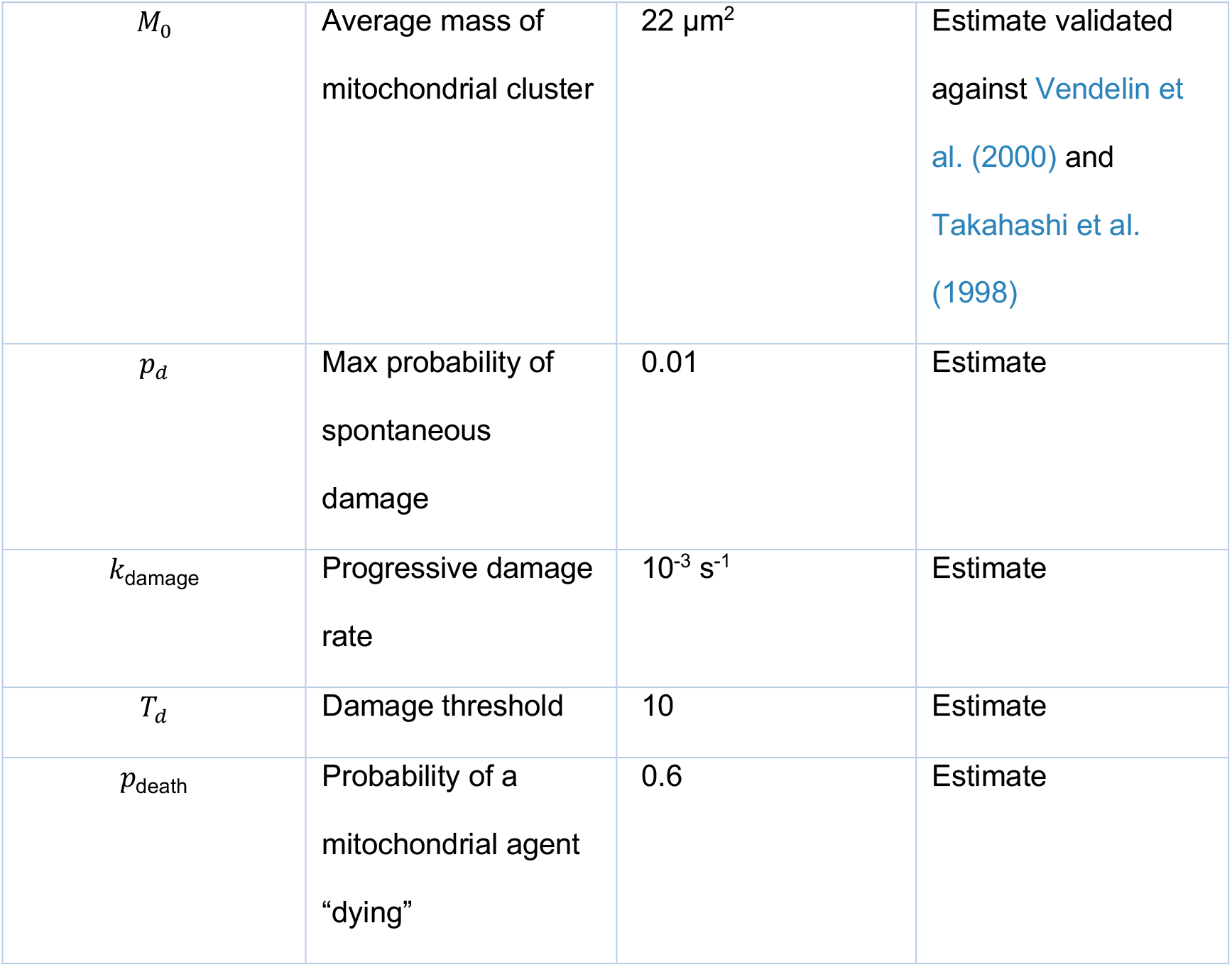
Parameter estimates for the PDE and ABM models

**Table 4.**
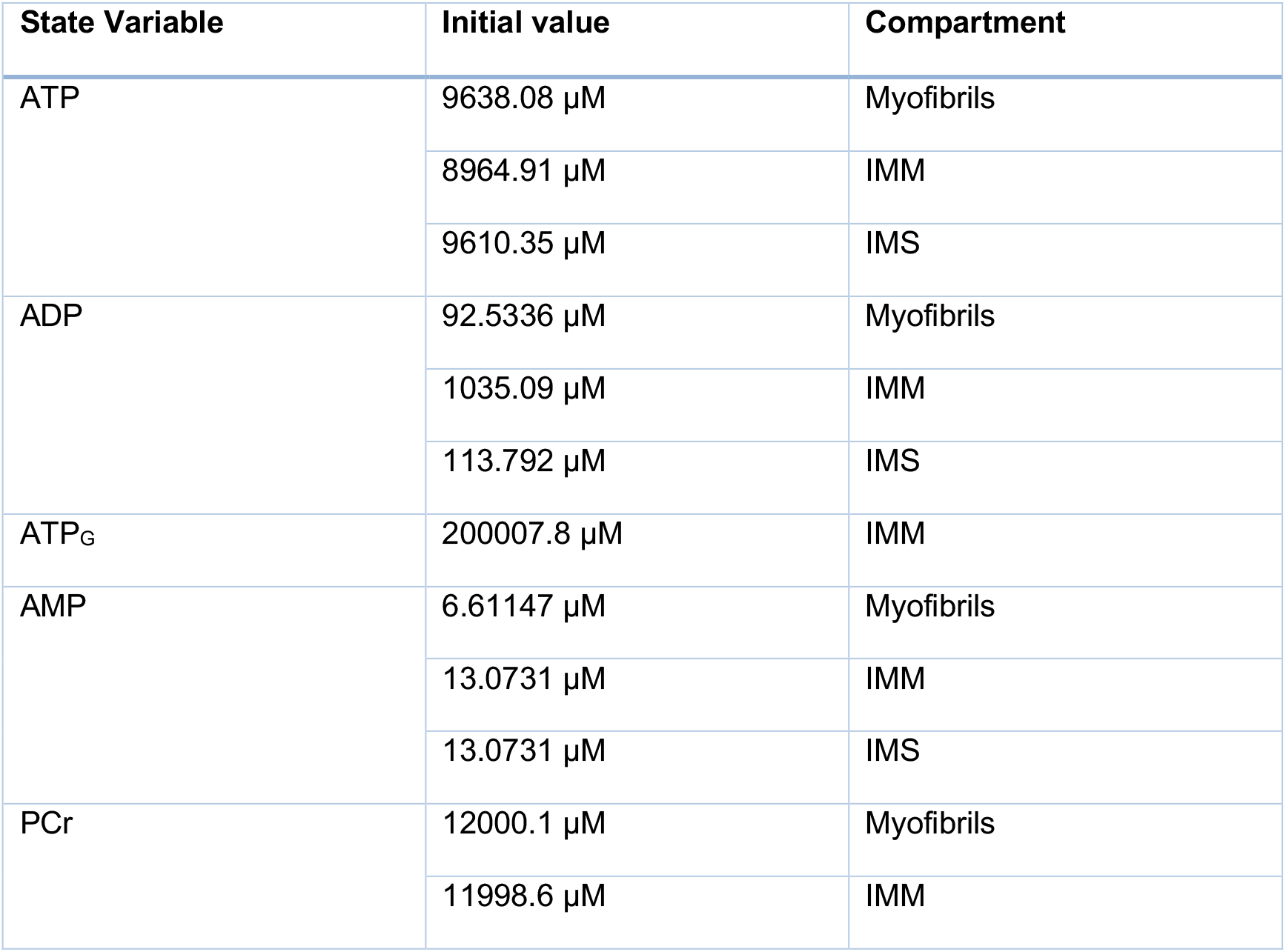
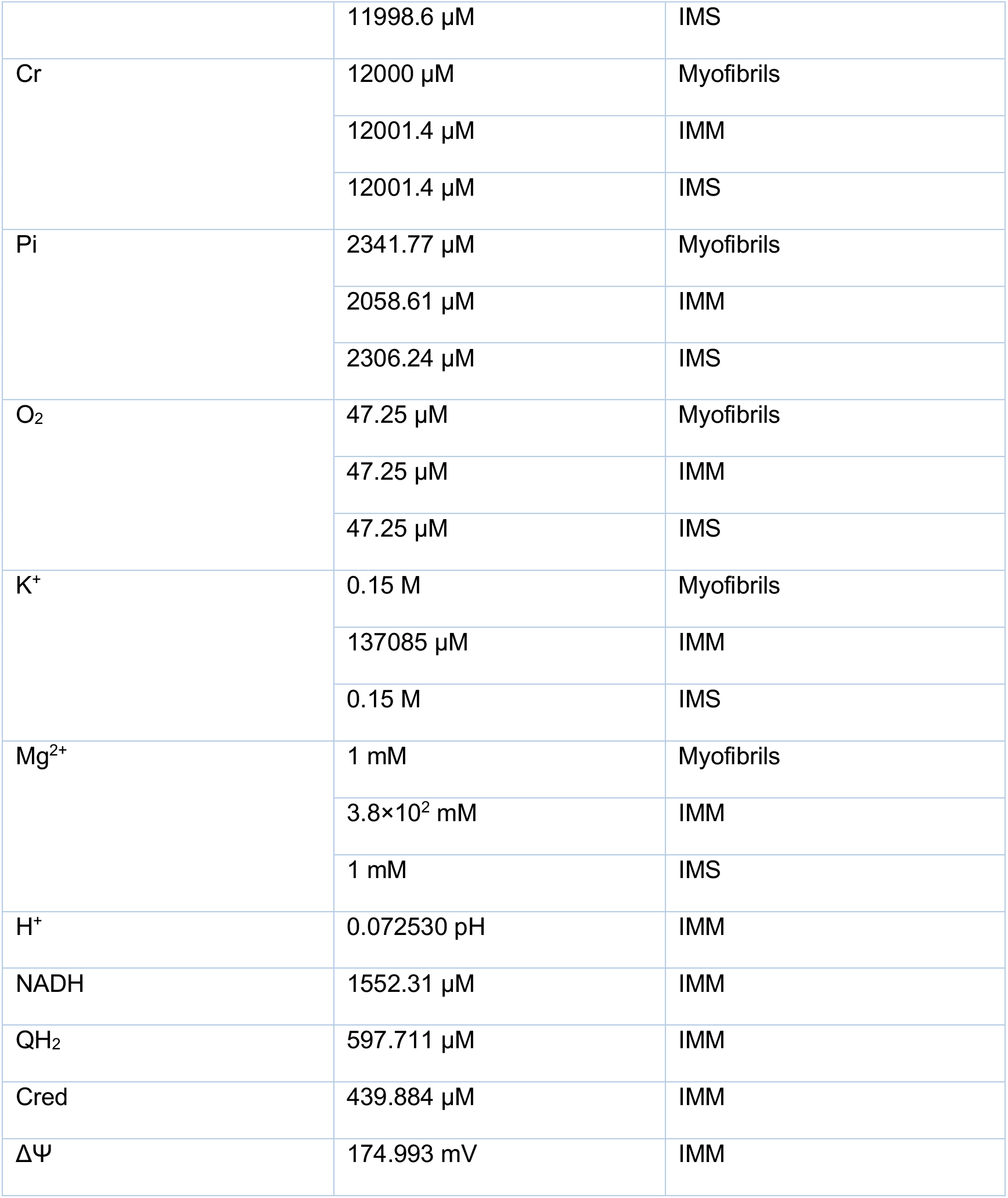
Initial conditions for PDE model

| State Variable | Initial value | Compartment |
| --- | --- | --- |
| ATP | 9638.08 $\mu\text{M}$ | Myofibrils |
| | 8964.91 $\mu\text{M}$ | IMM |
| | 9610.35 $\mu\text{M}$ | IMS |
| ADP | 92.5336 $\mu\text{M}$ | Myofibrils |
| | 1035.09 $\mu\text{M}$ | IMM |
| | 113.792 $\mu\text{M}$ | IMS |
| ATP<sub>G</sub> | 200007.8 $\mu\text{M}$ | IMM |
| AMP | 6.61147 $\mu\text{M}$ | Myofibrils |
| | 13.0731 $\mu\text{M}$ | IMM |
| | 13.0731 $\mu\text{M}$ | IMS |
| PCr | 12000.1 $\mu\text{M}$ | Myofibrils |
| | 11998.6 $\mu\text{M}$ | IMM |
| | 11998.6 $\mu\text{M}$ | IMS |
| Cr | 12000 $\mu\text{M}$ | Myofibrils |
| | 12001.4 $\mu\text{M}$ | IMM |
| | 12001.4 $\mu\text{M}$ | IMS |
| Pi | 2341.77 $\mu\text{M}$ | Myofibrils |
| | 2058.61 $\mu\text{M}$ | IMM |
| | 2306.24 $\mu\text{M}$ | IMS |
| O<sub>2</sub> | 47.25 $\mu\text{M}$ | Myofibrils |
| | 47.25 $\mu\text{M}$ | IMM |
| | 47.25 $\mu\text{M}$ | IMS |
| K<sup>+</sup> | 0.15 M | Myofibrils |
| | 137085 $\mu\text{M}$ | IMM |
|  | 0.15 M | IMS |
| Mg<sup>2+</sup> | 1 mM | Myofibrils |
| | 3.8$\times 10^2$ mM | IMM |
|  | 1 mM | IMS |
| H<sup>+</sup> | 0.072530 pH | IMM |
| NADH | 1552.31 $\mu\text{M}$ | IMM |
| QH<sub>2</sub> | 597.711 $\mu\text{M}$ | IMM |
| Cred | 439.884 $\mu\text{M}$ | IMM |
| $\Delta\Psi$ | 174.993 mV | IMM |

**Table 5.** Very large changes to enzyme activity with basal fission and fusion rates

| Fold change in complex I | Fold change in complex III | Fold change in complex IV | Fold change in complex V | ADP/ATP ratio ($\times 10^3$) | $\Delta\Psi$ | Mitochondrial volume fraction | Median cluster size |
| --- | --- | --- | --- | --- | --- | --- | --- |
| 1 | 1 | 1 | 1 | 9.1677 | 163.75 | 0.37539 | 9.24 |
| 10<sup>-4</sup> | 10<sup>-3</sup> | 10<sup>-3</sup> | 10<sup>-2</sup> | 53.68 | 125.69 | 0.41369 | 19.305 |
| 10<sup>-4</sup> | 10<sup>-2</sup> | 10<sup>-2</sup> | 10<sup>-3</sup> | 30.17 | 135.76 | 0.40281 | 16.83 |
| 10<sup>-3</sup> | 10<sup>-4</sup> | 10<sup>-4</sup> | 10<sup>-2</sup> | 115.67 | 119.65 | 0.43929 | 22.44 |
| 10<sup>-3</sup> | 10<sup>-2</sup> | 10<sup>-2</sup> | 10<sup>-4</sup> | 37.96 | 147.12 | 0.40304 | 16.17 |
| $10^{-2}$ | $10^{-4}$ | $10^{-4}$ | $10^{-3}$ | 98.13 | 118.42 | 0.42791 | 19.8 |

Finally, previous modelling work conducted by Dalmasso et al. (2017) suggests that mitochondrial populations establish and maintain homoeostasis, not by fission and fusion, but rather by mitochondrial motility. However, in cardiomyocytes mitochondria are organized into parallel columns extending along the length of the cell, which impairs motility (Cao and Zheng, 2019). Specifically, Eisner et al. (2017) observe that mitochondria in cardiomyocytes do not exhibit motility in vivo. However, the impact of cross-sectional network morphology on bioenergetics is still unresolved in our two-dimensional model. Additionally, our model does not necessarily distinguish between increased connectivity due to increased tethering of inter-mitochondrial junctions (IMJ) versus increased connectivity due to mitochondrial fusion. For example, Picard et al. (2013) note that acute exercise increases both IMJ tethering and mitochondrial mass, without an increase in fission or fusion (as quantified by the expression of fusion proteins such as Mfn1, Mfn2, and Opa1, and fission proteins such as Drp1 and Fis1). Future work will address this by using a finite element method to generalize the model to three dimensions, and then metabolically coupling it to an experimentally validated model of IMJ coupling that we intend to develop.

### Our model provides several experimentally testable predictions

Firstly, our model simulations predict that changes in fission and fusion activity – at least on a timescale of two minutes – do not <u>substantially</u> affect bioenergetics, namely, ΔΨ. Additionally, our model predicts that as workload, quantified via VO_2_, increases, so too does the frequency of fission and fusion events across the cell, and the median mitochondrial cluster size. While hypoxia is generally understood to induce mitochondrial fragmentation (when studied *in vitro* on a timescale of hours), our simulations suggest that this is preceded by a brief moment of mitochondrial aggregation. In other words, during hypoxia, mitochondria may acutely, i.e., on a timescale of two minutes, aggregate together before fragmenting or undergoing elongation. We emphasize that as simulations from a mathematical model, our results are hypothetical and as such, highlight the need for systematic quantitative measurements of mitochondrial dynamics. Indeed, we will refine our model along with our model assumptions as more experimental data becomes available.

### Our model highlights the need for quantitative, mechanistic understanding of mitochondrial dynamics to identify pathways for novel therapies

For example, supposing fission and fusion events do modulate ETC activity in cardiomyocytes at basal conditions, then how large a change in ΔΨ – either directly or via changes in enzyme activity in the ETC – do we observe? Moreover, our model assumes that we can induce a set change in fission and fusion activity. Experimentally, however, this is challenging in part because mitochondria can change their shape without necessarily increasing their expression of fission and fusion proteins (Picard et al., 2013). This leads to an additional question – can we induce changes in fission and fusion activity in a manner that is decoupled from inducing deleterious change in cell function (e.g., hypoxia or UV-induced damage). And finally, when and under what circumstances do mitochondria “switch” from fusion-dominated dynamics to fission-dominated dynamics, to minimize cellular stress. These experiments will provide critical insights into how mitochondrial form and cardiac metabolism are linked, and as a consequence help either robustly validate or negate our model’s findings with solid quantitative data.

In conclusion, our model suggests that ATP synthesis is robust to changes in fission and fusion rates. By combining experimental data with a system of mathematical equations, we developed a model that accounts for what has been speculated in the literature. We demonstrated that mitochondria achieve this robust adaptability by dynamically upregulating the number of fission and fusion events using a simple feedback-feedforward mechanism. Our modelling results suggest that changes in ATP synthesis might stem from changes to the respiratory-chain machinery caused by fission or fusion events. Indeed, our study leads to an interesting question, if in both healthy and damaged cardiomyocytes the ADP/ATP ratio is robust to changes in fission and fusion, when and under what circumstances are bioenergetics impacted?

## ACKNOWLEDGMENTS

The authors would like to thank Siavash B. Kalkhoran and Derek J. Hausenloy for their critical reading of this manuscript. A.K. was supported by an Australian Government Research Training Program (RTP) Scholarship; P.S.K. was supported by an Australian Research Council Discovery Project [DP180101512]; and S.G. and V.R. were supported by an Australian Research Council Discovery Project [DP170101358].

## AUTHOR CONTRIBUTIONS

Conceptualization, A.K., S.G, P.S.K., and V.R.; Methodology, A.K., S.G., and V.R.; Software, A.K. and S.G.; Formal Analysis, A.K and S.G.; Writing – Original Draft, A.K.; Writing – Review & Editing, A.K., S.G., P.S.K, and V.R; Supervision, P.S.K. and V.R.

## METHODS

### Lead contact and materials availability

Further information and requests for resources and reagents should be directed to and will be fulfilled by the Lead Contact, Vijay Rajagopal.

### Experimental model and subject details

#### Animals

The initial geometry of the model was inspired from an image of a longitudinal section of the cell. This image was acquired as part of a three-dimensional stack of electron microscopy images of a block of cardiac tissue from the left ventricular wall of an adult male Sprague Dawley rat. Details of the tissue preparation and imaging protocol used to collect these images can be found in Hussain et al. (2018). The longitudinal image was subsequently processed to identify and demarcate mitochondria boundaries and subsequently used to initiate the simulations. All animal procedures followed guidelines approved by the University of Auckland Animal Ethics Committee (for animal procedures conducted in Auckland, Application Number R826).

### Method Details

To quantify the role of mitochondrial network morphology on bioenergetics, we formulate a hybrid PDE-ABM system. We model biochemical reactions with a system of experimentally validated reaction-diffusion equations on a rectangular domain [0, *L*] × [0, *H*]; and use an agent-based model to describe changes in mitochondrial network morphology such as fission, fusion and biogenesis. We assume a constant pH of 7.1 and unless stated otherwise all fluxes are functions of state variables.

#### Partial differential equation model

##### ATP consumption

To model bioenergetics in the myofibrillar region of the cell, we slightly modify the bioenergetic model of Ghosh (2019) who considers several populations: [ATP], [ADP], and [AMP], the concentration of adenosine triphosphate, adenosine diphosphate, and adenosine monophosphate; [Pi], the concentration of inorganic phosphate; [Cr] and [PCr], the concentration of creatine and phosphocreatine; [O_2_], the concentration of oxygen; and [MgATP] and [MgADP], the concentration of magnesium-bound ATP and ADP. A table of all state variables is provided in Table 1. The interactions between these populations are described with a PDE system:

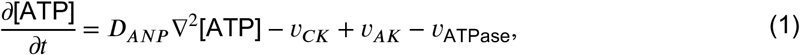

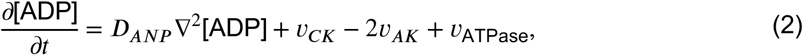

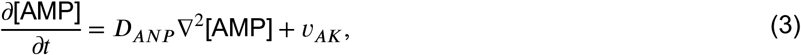

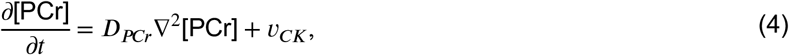

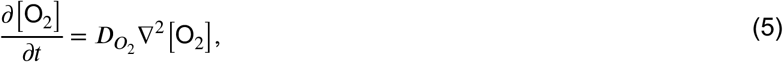

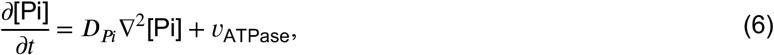

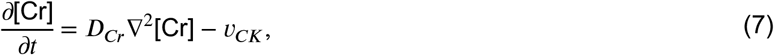

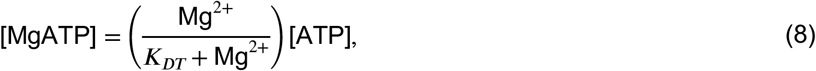

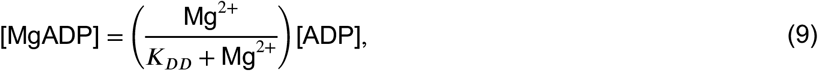

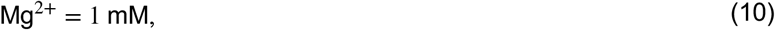

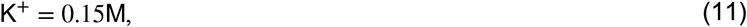

where the transport of metabolites across the cell is modelled using diffusion. Here, ATP is hydrolyzed – or consumed – at rate *υ*_ATPase_; ATP and AMP are catalyzed via adenylate kinase at rate *υ*_AK_; and creatine is converted into phosphocreatine via the creatine phosphate shuttle at rate *υ*_*ck*_. Details of these rates are provided in Table 2.

To approximate the cardiac cycle, we modify the ATP consumption rate used by Ghosh et al. (2018)

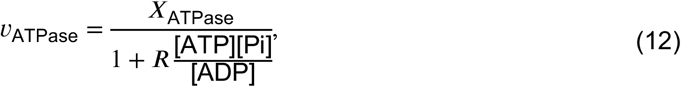

by multiplying it with a tent function 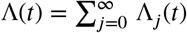, where

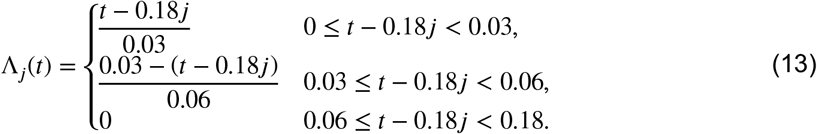

This function increases linearly from zero to one during the first 30 ms, decreases linearly to zero during the next 30 ms, and remains at zero until the end of the cardiac cycle at 180 ms. Here, *X*_ATPase_ is a model parameter that quantifies ATP consumption at various workloads and *R* is a fixed mass-action ratio. Unless stated otherwise, we assume a high-intensity workload of VO_2_=100 μmol min^-1^ g dw^-1^ corresponding to a value of *X*_ATPase_ = 5×10^4^ μM/s.

##### ATP production via OXPHOS

In Ghosh et al. (2018), the dynamics inside a mitochondrial matrix are described by two separate but metabolically linked PDE systems. One PDE system models the production of metabolites via OXPHOS in the inner mitochondrial membrane (IMM), while the other system models the transport of these metabolites from the IMM to the inter-membrane space (IMS). Once in the IMS, metabolites may diffuse into the myofibrillar region. To link these bioenergetic models to the ABM, we modify the OXPHOS model so that ETC enzyme activity and proton leakage depend on mitochondrial connectivity.

The production of ATP via OXPHOS in the IMM is described by the following system of PDEs:

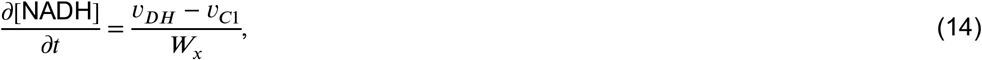

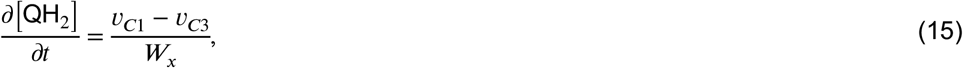

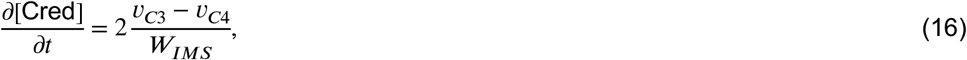

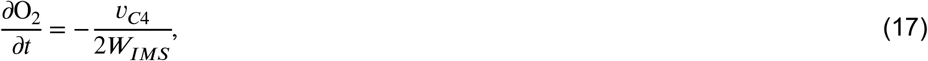

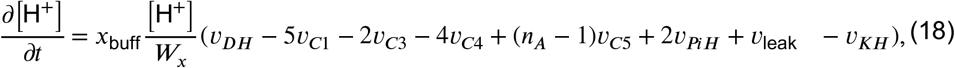

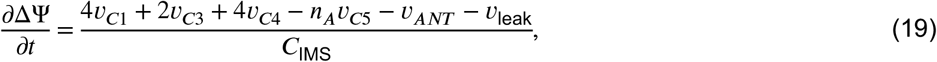

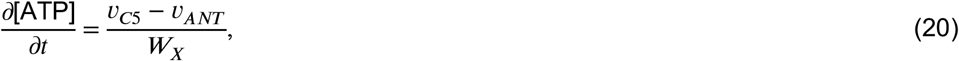

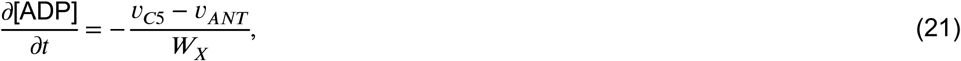

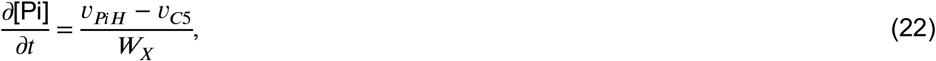

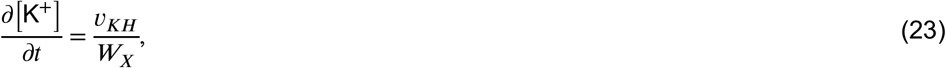

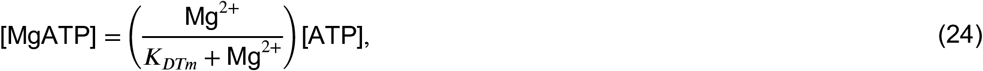

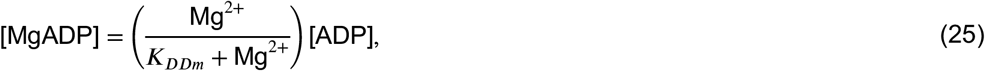

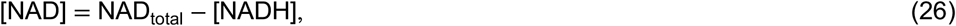

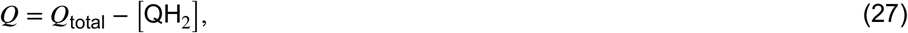

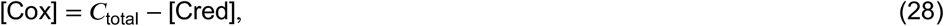

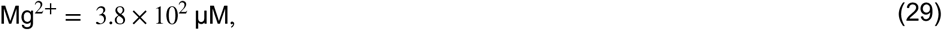

Here, ATP is produced via a series of protein complexes: complex I, complex III, complex IV and complex V at rates *υ*_*C*1_, *υ*_*C*3_, *υ*_*C*4_, and *υ*_*C*5_. While it is known that mitochondrial connectivity affects the electron transport chain (Fu et al., 2019; Parra et al., 2011; Pernas and Scorrano, 2016; Youle and van der Bliek, 2012), it is unclear if all or only some complexes are affected. Accordingly, we multiply the rates *υ*_*C*1_, *υ*_*C*3_, *υ*_*C*4_, and *υ*_*C*5_ by

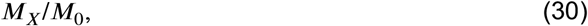

where *M*_*X*_ is the mitochondrial mass of a given matrix and *M*_0_ is the average mass of a mitochondrial matrix – thereby assuming that mitochondrial connectivity increases protein complex activity.

Mitochondrial connectivity may also modulate proton leakage (Fu et al., 2019; Parra et al., 2011; Pernas and Scorrano, 2016; Youle and van der Bliek, 2012). Moreover, mitochondrial damage can depolarize membrane potentials via increased proton leakage (Halestrap et al., 2004; Matsuda et al., 2010; Zorov et al., 2014; Zorova et al., 2018; Park et al., 2011). We account for these observations by multiplying the rate of proton leakage *υ*_leak_, by

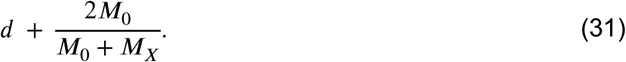

The first term in Equation 31 ensures that proton leakage increases with mitochondrial damage, *d*, while the second term,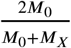, ensures that mitochondrial connectivity *decreases* proton leakage.

Additionally, ATP is transported to the IMS via adenine nucleotide translocase (ANT) at rate *υ*_*ANT*_; inorganic phosphate is co-transported at rate *υ*_*PiH*_; and potassium and protons are exchanged at rate *υ*_*KH*_. Finally, dehydrogenase flux stemming from the citric acid cycle occurs at rate *υ*_*DH*_.

The transport of metabolites in the IMM to the IMS is described by the following system of PDEs:

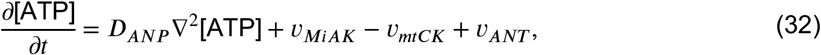

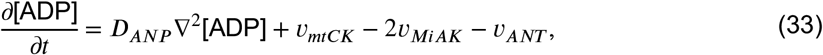

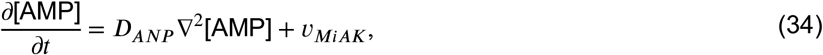

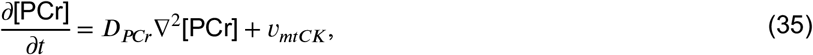

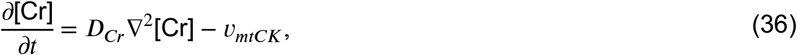

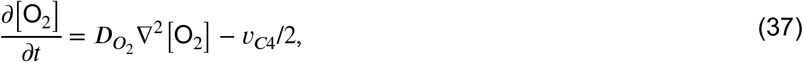

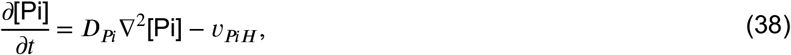

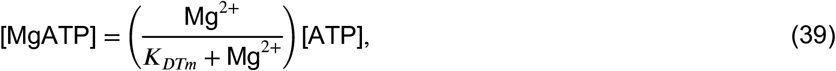

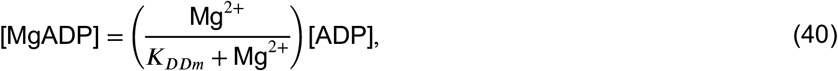

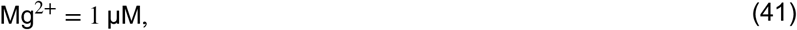

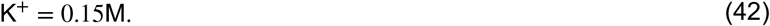

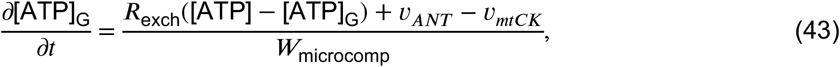

Here, ATP is transported via the protein adenine nucleotide translocase at rate *υ*_*ANT*_; ATP and AMP in the IMM are catalyzed via mitochondrial adenylate kinase at rate *υ*_*MiAK*_; creatine is converted into phosphocreatine in the IMM at rate *υ*_*mtCK*_; inorganic phosphate is co-transported at rate *υ*_*PiH*_; and oxygen is consumed in the IMM at rate *υ*_*C*4_/2. Equation 43, is a microcompartment between ANT and mitochondrial CK, that based on previous work by Aliev and Saks (1997), models phosphocreatine shuttling. Details of these rates are provided in Table 2.

##### Boundary conditions and initial conditions

As implemented by Ghosh et al. (2018), we impose no-flux boundary conditions (BC) on all state variables except oxygen, for which we impose a constant Dirichlet BC of 47.25 μM on the boundary. We use constant initial conditions, with details provided in Table 4.

#### Agent based model

Increased ATP demand along with oxidative stress is conducive to mitochondrial fusion and biogenesis, along with fission of damaged mitochondria (Dalmasso et al., 2017; Mihaylova and Shaw, 2011; Toyama et al., 2016; Egan et al., 2011). To this end, we model changes in network morphology with an agent-based model.

### Energetic stress

To model biophysical stressors, we introduce the concept of energetic stress

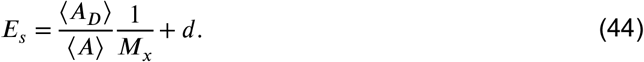

The fraction ⟨*A*_*D*_⟩/⟨*A*⟩ is the average ADP to ATP ratio within the mitochondrial matrix, acting as a measure of biophysical stress. Parra et al. (2011) speculate that increased connectivity improves bioenergetics by more uniformly distributing the mitochondrial membrane potential. We account for this by introducing a connectivity penalty to energetic stress of the form 1/*M*_*X*_, where *M*_*X*_ denotes the mitochondrial mass of a given matrix. The final term *d* denotes the level of mitochondrial damage (see Damage)

### Fusion

We assume that the probability of a mitochondrion undergoing fusion at each time step is given by

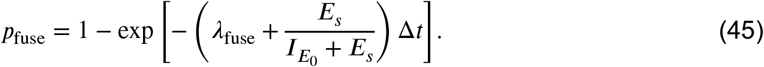

The size of each time step is denoted by Δ*t*, and the characteristic fusion rate is denoted by λ_fuse_. Our characterisation ensures that as energetic stress, *E*_*s*_, increases, the probability of fusion also increases. Here, 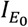 is a stress saturation constant. If a fusion event occurs, our mitochondrion (hereafter referred to as an agent) will fuse with all adjacent agents, *unless* the mass of the resultant agent exceeds *M*_max_.

### Biogenesis

Similarly, we assume the probability of a mitochondrion undergoing biogenesis at each time step is given by

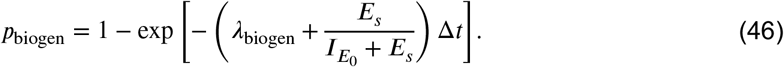

The parameter *λ*_biogen_ denotes the characteristic biogenesis rate. If a biogenesis event occurs, our mitochondrial matrix increases its mass by a single mitochondrion, *unless* the mass of the resultant matrix exceeds *M*_max_ or if there is no free space. The biogenesis process is implemented by associating a vacant cell to either the left or right of the current mitochondrial matrix with our agent. We assume that the contents of the cytosol are pushed away and distributed equally amongst neighboring cells.

### Fission

We assume that the probability of a mitochondrion undergoing fission at each time step is given by

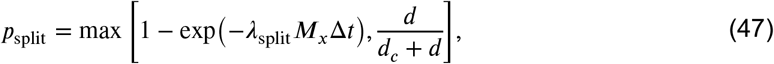

where *λ*_split_ is the characteristic fission rate and *M*_*x*_ the mitochondrial mass of a given matrix. The level of mitochondrial damage is described by *d* ≥ 1, and the extent to which mitochondrial damage drives fission is described by *d*_*C*_.

Our characterization assumes that in healthy mitochondria, fission occurs independently of energetic stress and is proportional to the number of agents in the given matrix. In Glancy et al. (2017), the authors note that damaged mitochondria rapidly increase fission to minimize the propagation of mitochondrial dysfunction. We capture this behavior by assuming that the probability of fission in damaged mitochondria is driven by damage according to the term 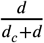 As local damage increases, the probability of fission approaches one. The switch between basal fission and damaged-induced fission occurs when the probability of damaged-induced fission matches the probability of basal-level fission, that is, when 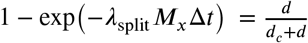 However, this switching condition is purely phenomenological and as such requires further experimentation to either be phenomenologically refined or replaced with aa mechanistic model.

Suppose our agent has an initial mitochondrial mass of *M*. Now let *M*_single_ denote the mass of a single mitochondrion. We assume that fission only occurs if *M* > 2*M*_single_. This implies that an agent must consist of more than two linked mitochondria for fission to occur, which is implemented for computational reasons. If fission occurs, the original agent divides into two new agents of mass *M*_1_ and *M*_2_, where *M*_1_ is chosen as a uniform random variable between *M*_single_ and *M* − *M*_single_ and *M*_2_ = *M* − *M*_1_.

### Damage

Mitochondria segregate damaged mitochondria via fission (Twig et al., 2008; Youle and van der Bliek, 2012; Glancy et al., 2017). Mitochondrial damage is described by the variable *d* and is assumed to exist in two states, low and high. We assume that after fission, the probability of the newly separated mitochondria becoming damaged is *p*_damage_. This probability is defined as

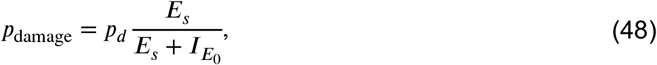

where the parameter *p*_*d*_ describes the maximal probability of mitochondrial damage. The factor 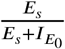 ensures that increased energetic stress results in an increased likelihood of mitochondrial damage.

Damaged mitochondria start from a low-damage state, corresponding to *d* = 1 and increases by 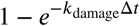 every time step. Once damage hits a critical threshold *d* = *T*_*d*_, our state switches from a low-damage state to a high-damage state. Mitochondria that are highly damaged are assumed to be susceptible to increased turnover (Hamacher-Brady and Brady, 2016). The probability of mitochondrial turnover is given by *p*_death_ > 0.5. This approach to modelling mitochondrial damage is similar to the approach utilised by Dalmasso et al. (2017). Here, turnover is not referring to cell death nor mitophagy *per se*, but rather refers to a mitochondrial agent dying.

Healthy mitochondria are treated as having a zero-damage state, i.e., *d* = 0. If two mitochondria with damage states of *d*_1_ and *d*_2_ fuse, then the resultant damage is assumed to be the average of the two, i.e. (*d*_1_ + *d*_2_)/2. If this value is below one, the mitochondria are no longer marked as damaged.

#### Parameter estimates

A summary of parameter values is provided in Table 3. Where possible, we have used data from animal models to characterize our estimates; however, some of the available data comes from *in vitro* models due to the limited availability of animal data. For the PDE model, we used the flux terms implemented by Ghosh et al. (2018). Details of how these parameters were estimated are provided therein. We used manual calibration instead of formal parameter fitting, which in our case is not feasible due to a lack of data directly corresponding to specific model parameters. We summarize how we obtained these estimates for our ABM parameters below. Unless stated otherwise, we assume a high-intensity workload of VO_2_=100 μmol min^-1^ g dw^-1^.

### Agent-based model

- *λ*_split_: As a plausible estimate, we use a characteristic fission rate of 2×10^−3^ s^-1^.
- *λ*_fuse_: Using a murine cardiomyocyte model, Eisner et al. (2017) estimate a characteristic fusion rate of 1.67×10^−2^ s^-1^.
- *λ*_biogen_: In Dalmasso et al. (2017) the authors optimize a mathematical model to estimate a characteristic biogenesis frequency of 28.9 minutes, which equates to 1/28.9 min^-1^ = 5.77×10^−4^ s^-1^.
- *M*_0_, *M*_single_, and *M*_max_: Based on our image data we estimate *M*_single_=8.25×10^−1^ μm^2^ and *M*_max_= 43 μm^2^. Motivated by this we set 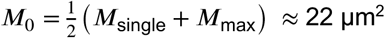.
- *d*_*C*_, *p*_*d*_, *k*_damage_, and *T*_*d*_: As plausible estimates, we use *d*_*C*_=250, *p*_*d*_=0.01,*k*_damage_=10^−3^ s^-1^, and *T*_*d*_=10.
- *p*_death_: We arbitrarily set the probability of a mitochondrial agent dying to be 0.6.

#### Simulations

The initial geometry of our mitochondrial network was inspired by tissue samples taken from a Sprague Dawley rat. Our initial conditions are a simplified 2D electron microscopy representation of a healthy rat heart and are visualized in Figure 1A. We assume that the lateral and longitudinal dimensions of our hypothetical cardiomyocyte are 79 μM and 15.75 μM respectively. Our spatial increments, *Δx* and *Δy* are taken to be 1 μM and 0.75 μM respectively. These increments coincide with the typical dimensions of a fibre-parallel mitochondrion, allowing us to model mitochondrial matrices as mesh-points on our domain. We note that using a different initial condition does not appear to change our results (see Figure S1).

Our PDEs are discretized using the method of lines and solved using semi-implicit Strang splitting. Our linear and non-linear components are both solved with MATLAB’s inbuilt stiff ODE solver “ode15s” with an absolute error tolerance of 10^−6^. Our ABM time step, *Δt*, is set to be 0.01 s.

To determine the minimum number of ABM runs, we screen for variance stability (Lee et al., 2015; Lorscheid et al., 2012), i.e., identify the number of runs required for the coefficient of variation to be less than some fixed tolerance. We find that with 5 runs, the coefficients of variation for our ADP/ATP ratios are below 10^−2^, which is considered acceptable in the literature (Lee et al., 2015). Thus, the results from our ABM represent an average from 5 runs unless stated otherwise.

## SUPPLEMENTAL INFORMATION

**Movie S1. Simulation of the model at baseline conditions**.

Inset depicts a moving histogram depicting the distribution of mitochondrial (normalized so that the total area sums to one). Also depicted are the ADP/ATP ratios; Pi concentrations; and the ΔΨ’s predicted by the model. These values oscillate for each cardiac cycle. Related to Figure 3.

**Movie S2. Simulation of the model under hypoxia**.

The value of O_2_ on the boundary is 5 μM at a workload of VO_2_ = 100 μmol min^-1^ g dw^-1^. Inset depicts a moving histogram depicting the distribution of mitochondrial (normalized so that the total area sums to one). Additionally depicted are the ADP/ATP ratios; O_2_ concentrations; and the ΔΨ’s predicted by the model. These values oscillate for each cardiac cycle. Related to Figure 4C.

**Figure S1.**
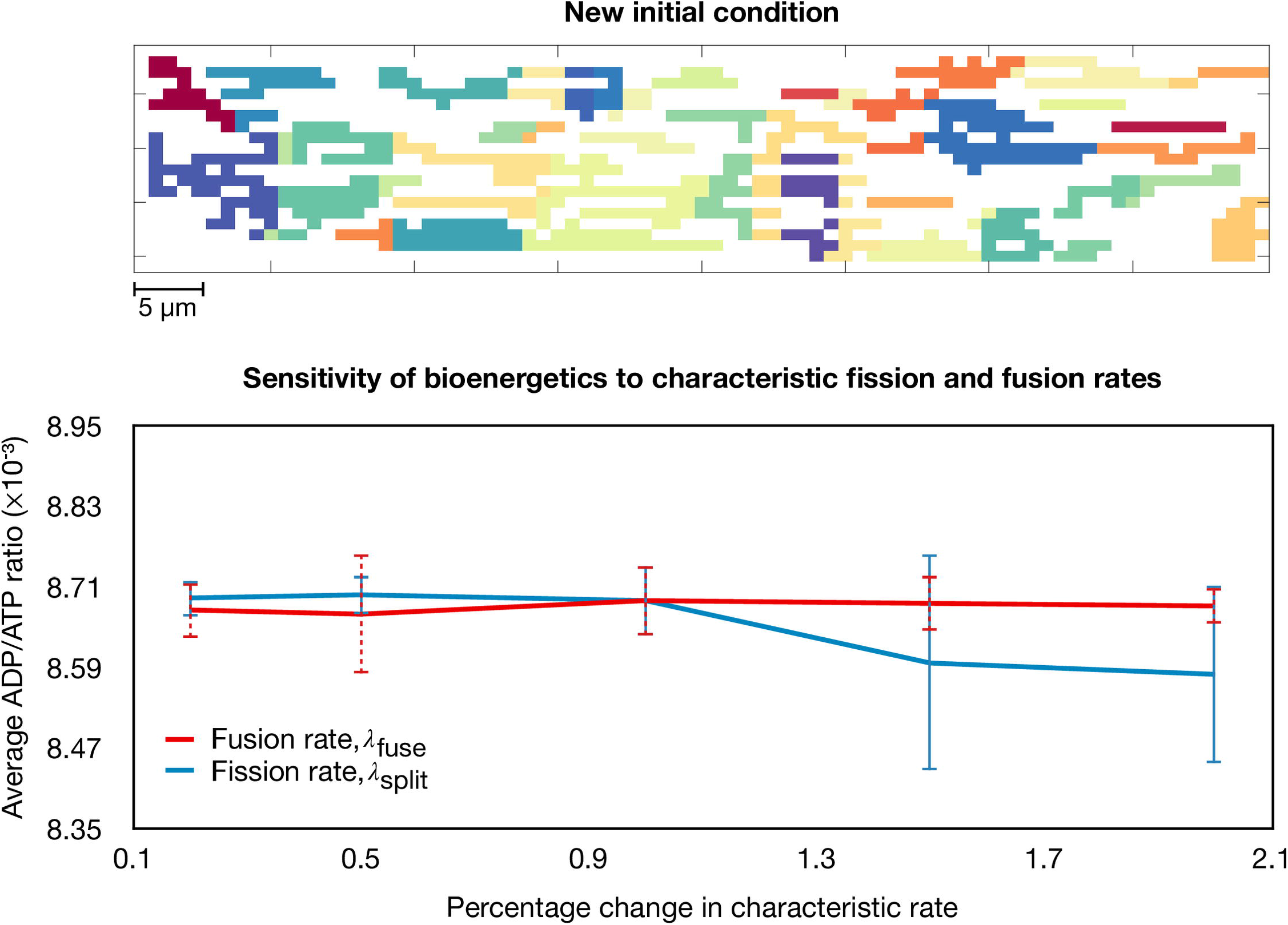
Sensitivity of ADP/ATP ratio with different initial conditions. Bioenergetics are robust to changes in the characteristic fission rate, *λ*_split_, and fusion rate, *λ*_split_, regardless of the initial condition used in the model. Related to Methods.

